# MAUI-seq: Metabarcoding using amplicons with unique molecular identifiers to improve error correction

**DOI:** 10.1101/538587

**Authors:** Bryden Fields, Sara Moeskjær, Ville-Petri Friman, Stig U. Andersen, J. Peter W. Young

## Abstract

**Background:** Sequencing and PCR errors are a major challenge when characterising genetic diversity using high-throughput amplicon sequencing (HTAS).

**Results:** We have developed a multiplexed HTAS method, MAUI-seq, which uses unique molecular identifiers (UMIs) to improve error correction by exploiting variation among sequences associated with a single UMI. We show that two main advantages of this approach are efficient elimination of chimeric and other erroneous reads, outperforming DADA2 and UNOISE3, and the ability to confidently recognise genuine alleles that are present at low abundance or resemble chimeras.

**Conclusions:** The method provides sensitive and flexible profiling of diversity and is readily adaptable to most HTAS applications, including microbial 16S rRNA profiling and metabarcoding of environmental DNA.

## Introduction

The evaluation of DNA diversity in environmental samples has become a pivotal approach in microbial ecology [1] and is increasingly also used to assess the distribution of larger organisms [2]. If a core gene can be amplified from environmental DNA with universal primers, the relative abundance of species in the community can be estimated from the proportions of species-specific variants among the amplicons. High throughput amplicon sequencing (HTAS), often termed metabarcoding, has become a cost-effective way to detect multiple species simultaneously within a range of environmental samples [3–8]. While shotgun sequencing of the whole community (metagenomics) can provide a richer description of the functions in a community, HTAS remains a more efficient tool for comparing the species diversity of a large number of community samples. Despite the extensive use of HTAS for interspecies ecological diversity studies, few investigations have utilised HTAS for intraspecies analysis [9, 10]. As 16S rRNA amplicons are too highly conserved to estimate microbial within-species diversity, other target gene candidates need to be considered in order to sufficiently discern intraspecies sequence variation.

Many studies have evaluated the extent of PCR-based amplification errors and bias for HTAS diversity studies [4, 6, 7, 11]. Numerous known PCR biases reduce the accuracy of diversity and abundance estimations, with the major concern being the inability to confidently distinguish PCR error from natural sequence variation in environmental samples, which is an especially limiting factor for intraspecific studies.

Polymerase errors, production of chimeric sequences by template switching, and the stochasticity of PCR amplification can be major causes of PCR errors [11–13]. Polymerase errors introduce new sequences into the template population during amplification. These sequence errors include not only substitutions but also insertions and deletions. The use of proofreading polymerases, optimised DNA template concentration, and reduced PCR cycle number have been suggested to reduce these errors [7, 11, 14].

In order to account for the introduction of sequence variants in PCR amplification, several sequence-classification approaches have been established to manage diversity estimates. The most common method is the use of operational taxonomic units (OTUs) in microbial diversity studies which analyse target gene sequences and cluster based on an arbitrary fixed similarity threshold (QIIME [15]; UPARSE [8, 16–20]. Within species boundaries this technique could dramatically reduce the resolution of naturally occurring sequence variation.

Most recent methods rely on the formation of sequence groups called amplicon sequence variants (ASVs) (DADA2, [19]; UNOISE3, [20, 21]. This approach allows sequence resolution down to one nucleotide, which is advantageous for determining intraspecies allelic variation, but noise from PCR errors is also more evident. Variation induced by PCR errors often cannot be differentiated from rare natural allelic variation without the use of sequence denoising methods [11]. DADA2 relies on a quality-aware parametric error model, which is developed on a per sequencing run basis. This increases the run time compared to UNOISE3, which uses a one-pass technique [22].

An approach that can reduce sequencing noise is to assign a unique molecular identifier (UMI) to every initial DNA template within an HTAS sample, which also enables evaluation of PCR amplification bias [23]. Additionally, the UMI provides a potential route to address polymerase errors in metabarcoding studies. The UMI is provided by a set of random bases in the gene-specific forward inner primer, which introduces a unique DNA sequence into every initial DNA template upstream of the amplicon region during the first round of amplification. Once all original DNA template strands are assigned a unique UMI, an outer forward primer and the gene-specific reverse primer can be used for further amplification. Consequently, all subsequent DNA amplified from the original template will have the same UMI, so the number of reads amplified from the initial template can be calculated. Grouping sequences by shared UMI allows identification of a consensus, which is assumed to be the correct sequence [24]. To our knowledge, UMIs have previously only been used for single-amplicon interspecies investigations [25–28].

Here, we present a method for metabarcoding using amplicons with unique molecular identifiers to improve error correction – MAUI-seq. The innovative approach is that we use variation among sequences associated with a single UMI to identify erroneous sequences, and we show that this improves error correction compared to non-UMI based analysis using the state-of-the-art software packages DADA2 and UNOISE3.

## Results

### Laboratory protocol: UMI labelling and amplicon multiplexing

We developed a procedure (MAUI-seq) to amplify multiple target genes from environmental samples, while assigning a random UMI to each initial copy of a template. We opted for a straightforward protocol using a “one-pot” initiation and amplification system. Forward primers consist of two modules; an inner primer bearing the UMI and designed to amplify the target gene, and a universal outer primer that binds only to a linker on the inner primer (**Figure 1A**). We used a 12-base UMI that allowed over 4 million distinct sequences, which is adequate to ensure that duplicate use is negligible for samples with a few thousand sequenced UMIs. For studies with greater sequencing depth, a longer UMI can easily be designed. As a test case, we used MAUI-seq to investigate the genetic diversity of the nitrogen-fixing bacterium *Rhizobium leguminosarum* symbiovar *trifolii (Rlt)* by characterising amplicons from the chromosomal core genes *rpoB* and *recA* and the plasmid-borne nodulation genes *nodA* and *nodD*. Each gene was amplified separately in a single reaction, using a target-specific inner forward primer (at low concentration) to assign the UMI and a universal outer primer (at high concentration) to amplify the resulting molecules (**Figure 1A**). The resulting amplicons were pooled and tagged by Nextera to identify the sample, then further pooled for high-throughput paired-end sequencing (**Figure 1B**). The full MAUI-seq step-by-step laboratory protocol can be found in **Additional File 1**.

**Figure 1.**
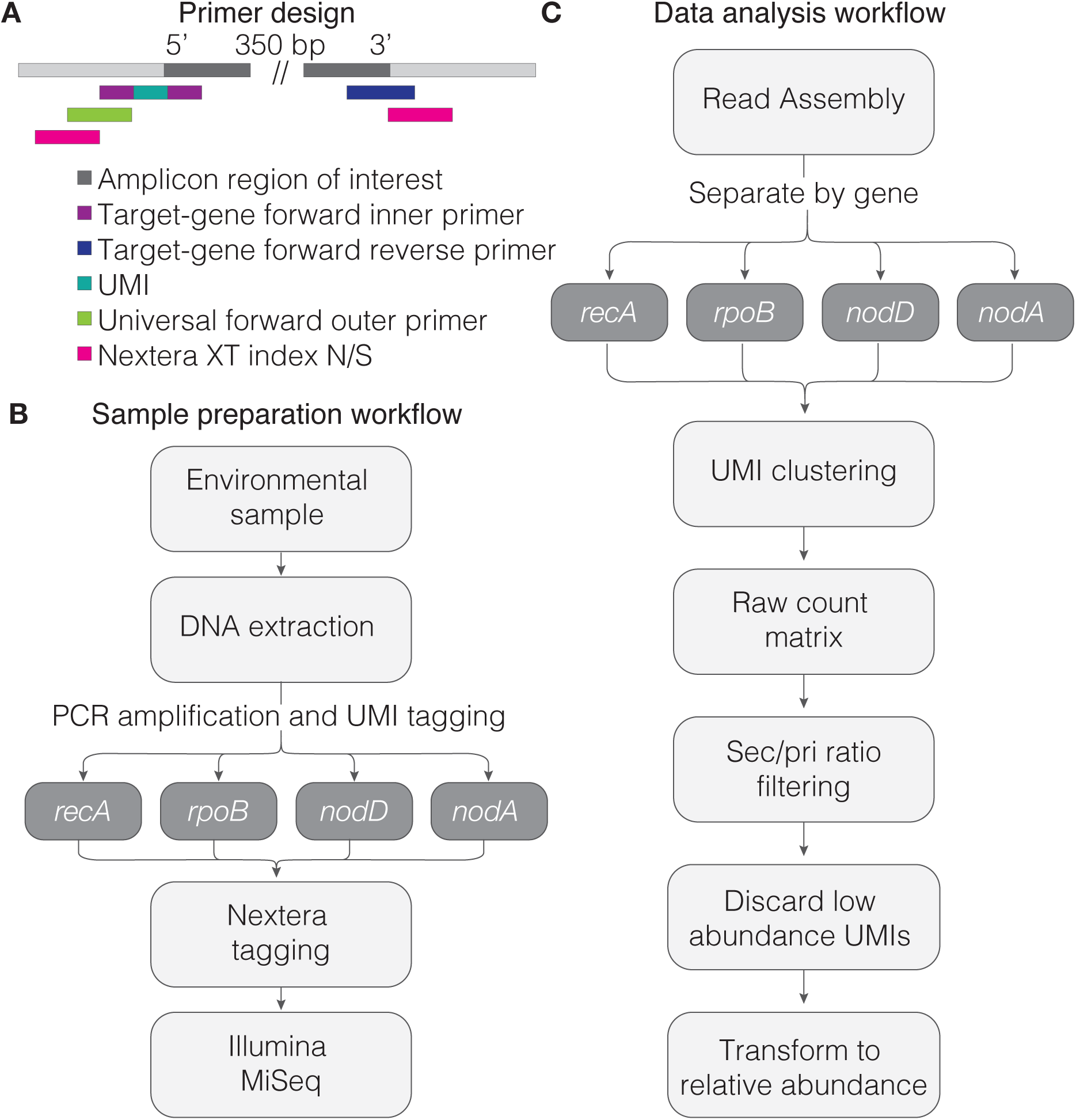
Primer design and method workflow. **A:** Primer design using the sense strand of the target DNA template as an example. The amplicon region of interest should be no longer than 500bp. The target-gene forward inner primer, universal forward outer primer and the target-gene reverse primer are all used in the initial PCR. The Nextera XT indices provide sample barcodes in a separate PCR step. The unique molecular identifier (UMI) region is shown in turquoise on the target-gene forward inner primer. **B:** Sample preparation workflow. **C:** MAUI-seq data analysis workflow.

### Analysis protocol: filtering using UMI-based error rates

The resulting paired-end reads were merged and then separated by gene prior to downstream analysis, where UMIs are critical in two ways. Firstly, sequences are clustered by UMI, and the number of unique UMIs is counted for each distinct sequence, selecting the most abundant sequence associated with each UMI (**Figure 1C**). UMIs are discarded as ambiguous if the most abundant sequence does not have at least two reads more than the next in abundance. The most abundant sequence will usually be the correct one (**Figure 2A** Case 1) but, because most UMIs are represented by just a small number of reads, it can sometimes happen that an erroneous sequence is sampled more often than the true sequence, so the primary sequence of the UMI becomes this erroneous sequence (**Figure 2A** Case 2). Secondly, we reasoned that it may be possible to eliminate these errors by using the UMIs to provide information on global error rates across all samples. We implemented this in MAUI-seq by noting both the most abundant (primary) and the second most abundant (secondary) sequence if two or more sequences were associated with the same UMI. MAUI-seq then distinguishes between true and erroneous sequences based on the ratio of primary and secondary occurrences of each sequence, eliminating sequences that show a high ratio (default is 0.7) of secondary to primary occurrences (**Figure 1C** and **Figure 2B**). The 0.7 threshold was chosen empirically, based on the ratios observed for known true and erroneous sequences, but it is a compromise because the incidence of secondary sequences varies across genes and studies. An examination of the results may suggest choosing different thresholds in other studies. Finally, globally rare sequences are discarded (default threshold is 0.1% averaged across samples - a lower threshold could be used if samples were sequenced to a greater depth). Python scripts for separating the genes and for the UMI analysis are available at https://github.com/jpwyoung/MAUI.

**Figure 2.**
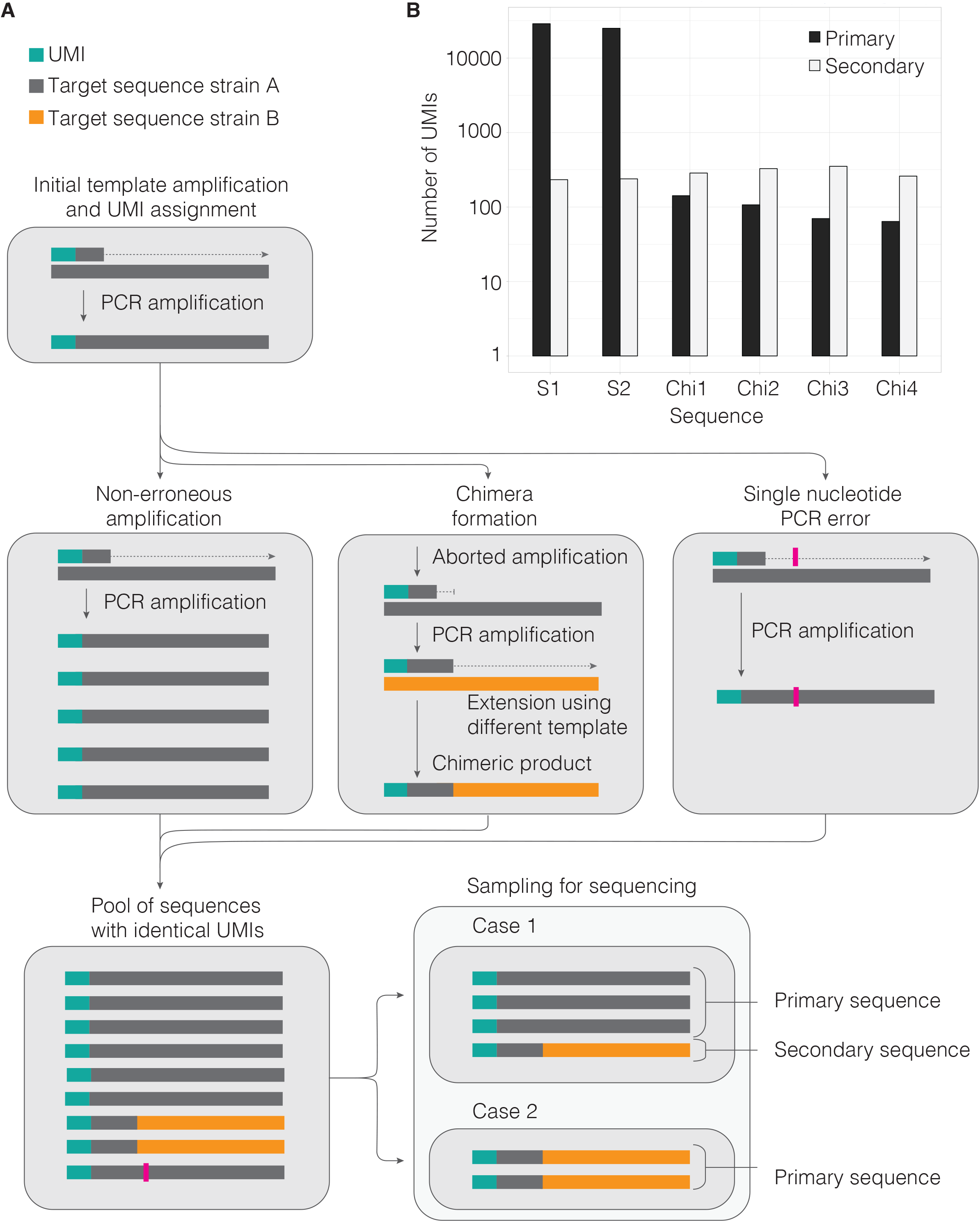
Erroneous read formation and filtering. **A:** Schematic showing the formation of different sequences with identical UMIs, and bias introduced when sampling for sequencing. **B:** Example data showing the occurrence of real and chimeric *rpoB* sequences as primary and secondary sequence (log scale). S1 and S2: Real sequences derived from two different rhizobium strains (SM170C and SM3). Chi1-4: Chimeric sequences.

### Validation using purified DNA mixed in known proportions

We first evaluated the accuracy of MAUI-seq by profiling DNA mixtures with known strain DNA ratios. DNA was extracted from two *Rlt* strains differing by a minimum of 3bp in each of their *recA*, *rpoB*, *nodA*, and *nodD* amplicon sequences, and the extracted DNA was mixed in different ratios (**Supplementary Table S1**). After amplification and sequencing, assembled reads were assigned to their target gene and analysed using MAUI-seq and two programs frequently used for de-noising of amplicon sequencing data, DADA2 and UNOISE3 [19, 21]. Since rare sequences have a high error rate, we discarded (for each of the three methods) sequences that fell below a threshold frequency of 0.1% of accepted sequences. The observed and expected strain ratios were highly correlated for all four genes across the three analysis methods, and we found that the performances of the proofreading (Phusion) and non-proofreading (Platinum) polymerases were gene-dependent, which could be due to differences in amplification efficiency for the four templates (**Table 1** and **Supplementary Figures S1-S4**). On average, MAUI-seq detected between 98.5% and 100% true sequences exactly matching those of the two strains in the mixture, while DADA2 ranged from 89.7% to 100%, and UNOISE3 from 79.8% to 100% (**Table 1**). The better performance of MAUI-seq was due to more effective elimination of chimeras, which were especially abundant when the PCR reaction was carried out using the Platinum non-proofreading polymerase (**Table 1** and **Supplementary Figures S1-S4**). For the proofreading polymerase, DADA2 detected 100% true sequences for all four genes, whereas MAUI-seq detected 99.03% for *nodA*, failing to eliminate three rare sequences that did not have sufficient secondary counts. This suggests that DADA2 performs equally well or even slightly better than MAUI-seq, when a proofreading polymerase is used to amplify DNA from a simple, two-component mix. The prevalence of secondary sequences varied with gene and polymerase: the secondary/primary ratio for accepted sequences was 0.0322 for *rpoB* using Phusion, but just 0.0002 for *nodD* using Platinum. When the ratio was very low, there were insufficient secondary counts for MAUI-seq to eliminate erroneous sequences effectively.

**Table 1.** Total number of detected sequences in the synthetic mix samples using MAUI-seq, DADA2 and UNOISE3. The percentage of true sequences is averaged over 23 samples for Platinum (non-proofreading) and 14 samples for Phusion (proofreading).

|  |  | Platinum |  |  | Phusion |  |  |  |
| --- | --- | --- | --- | --- | --- | --- | --- | --- |
|  |  | MAUI-seq | DADA2 | UNOISE3 | MAUI-seq | DADA2 | UNOISE3 | exp. seq* |
| *rpoB* | **n seq** | 2 | 3 | 4 | 2 | 2 | 2 | 2 |
|  | **%true\*** | 100 | 96.96 | 93.80 | 100 | 100 | 100 | - |
|  | **Cor.exp/obs\*** | 0.956 | 0.977 | 0.981 | 0.996 | 0.999 | 0.9998 | - |
|  | **chim.freq\*** | 0 | 0.07 | 0.13 | 0 | 0 | 0 | - |
| *recA* | **n seq** | 2 | 2 | 2 | 2 | 2 | 2 | 2 |
|  | **%true** | 100 | 100 | 100 | 100 | 100 | 100 | - |
|  | **Cor.exp/obs** | 0.984 | 0.991 | 0.989 | 0.948 | 0.952 | 0.947 | - |
|  | **chim.freq** | 0 | 0 | 0 | 0 | 0 | 0 | - |
| *nodA* | **n seq** | 6 | 5 | 4 | 5 | 2 | 4 | 2 |
|  | **%true** | 99.04 | 89.70 | 89.93 | 99.03 | 100 | 90.43 | - |
|  | **Cor.exp/obs** | 0.985 | 0.998 | 0.999 | 0.989 | 0.999 | 0.999 | - |
|  | **chim.freq** | 0.10 | 0.25 | 0.22 | 0.04 | 0 | 0.16 | - |
| *nodD* | **n seq** | 7 | 6 | 21 | 3 | 3 | 14 | 3<sup>†</sup> |
|  | **%true** | 98.49 | 93.93 | 90.10 | 100 | 100 | 79.83 | - |
|  | **Cor.exp/obs** | 0.998 | 0.998 | 0.995 | 0.990 | 0.998 | 0.995 | - |
|  | **chim.freq** | 0.05 | 0.05 | 0.13 | 0 | 0 | 0.11 | - |
| **all** | **%true-overall\*** | 99.76 | 93.73 | 91.93 | 99.74 | 100 | 91.71 | - |

\***n seq** is the total number of sequences occurring across all samples. **%true** is calculated by dividing the number of counts for the true sequences by the total number of counts accepted by the method. **%true-overall** is based on summed counts for all four genes. **Cor.exp/obs** is the Pearson correlation for the observed proportion of SM170C reads versus the expected proportion. **Chim.freq** is the proportion of chimeras compared to total reads at 0.5 expected proportion of sequences. **Exp.seq** is the expected number of detected sequences.
<sup>†</sup> SM170C has a second copy of *nodD* [29].

### Validation using environmental samples

To test the method on more complex samples, we compared *Rlt* populations in root nodules from two locations in Denmark, a clover trial station in Store Heddinge on Zealand and a lawn at Aarhus University in Jutland (the Field-Samples-1 dataset; **Supplementary Figure S5**). One hundred nodules were pooled for each sample and each plot was sampled in four replicates. Platinum Taq polymerase enzyme was used for amplification. Each clover root nodule is usually colonised by a single *Rhizobium* strain, so a maximum of 100 unique sequences per gene is expected per sample.

For Field-Samples-1, the total number of distinct sequences for MAUI-seq and DADA2 were in the same range as the number of distinct alleles observed in a population of 196 natural European *Rlt* isolates [29] (**Table 2**). In contrast, UNOISE3 produced a substantially higher number of distinct sequences, suggesting that its default filtering might be too lenient for our data (**Table 2**). The sequences accepted as true by MAUI-seq were nearly all also included in the DADA2 and UNOISE3 outputs (**Figure 3**). On the other hand, DADA2 and UNOISE3 both accepted a number of sequences that were filtered out by MAUI-seq, and many of these were eliminated by MAUI-seq because a high ratio of secondary to primary occurrences strongly suggested that they represent errors and not real sequences (**Figure 3** and **Additional file 2**). To provide independent evidence as to whether sequences were likely to be genuine, we checked whether they matched (or differed by a single nucleotide from) known sequences in either a reference database of 196 natural European *Rlt* isolates [29], or the NCBI whole-genome shotgun database (**Figure 3**). The great majority of sequences rejected by MAUI-seq did not have exact matches to these known sequences. A few sequences that exactly matched known alleles were included by DADA2 and UNOISE, but not by MAUI-seq. These sequences were not reported by MAUI-seq because their UMI counts were below the abundance threshold, not because the secondary/primary occurrence filter identified them as erroneous (**Figure 3**). The count threshold could be lowered to include rarer sequences, if the study required it.

**Table 2.**
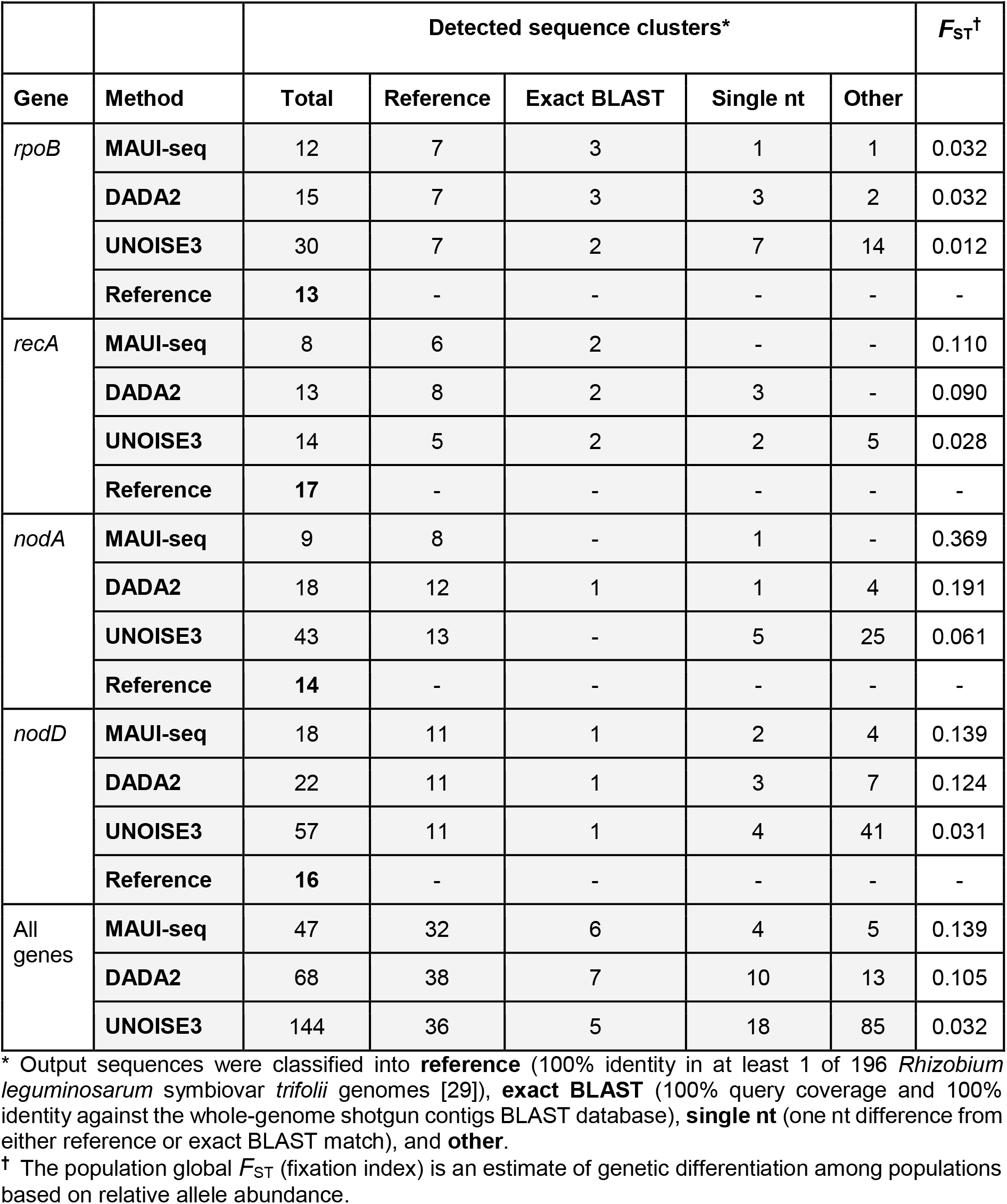
Total number of detected sequence clusters in root nodule samples (Field-Samples-1) using MAUI-seq, DADA2, and UNOISE3 clustering and genetic differentiation between populations.

**Figure 3.**
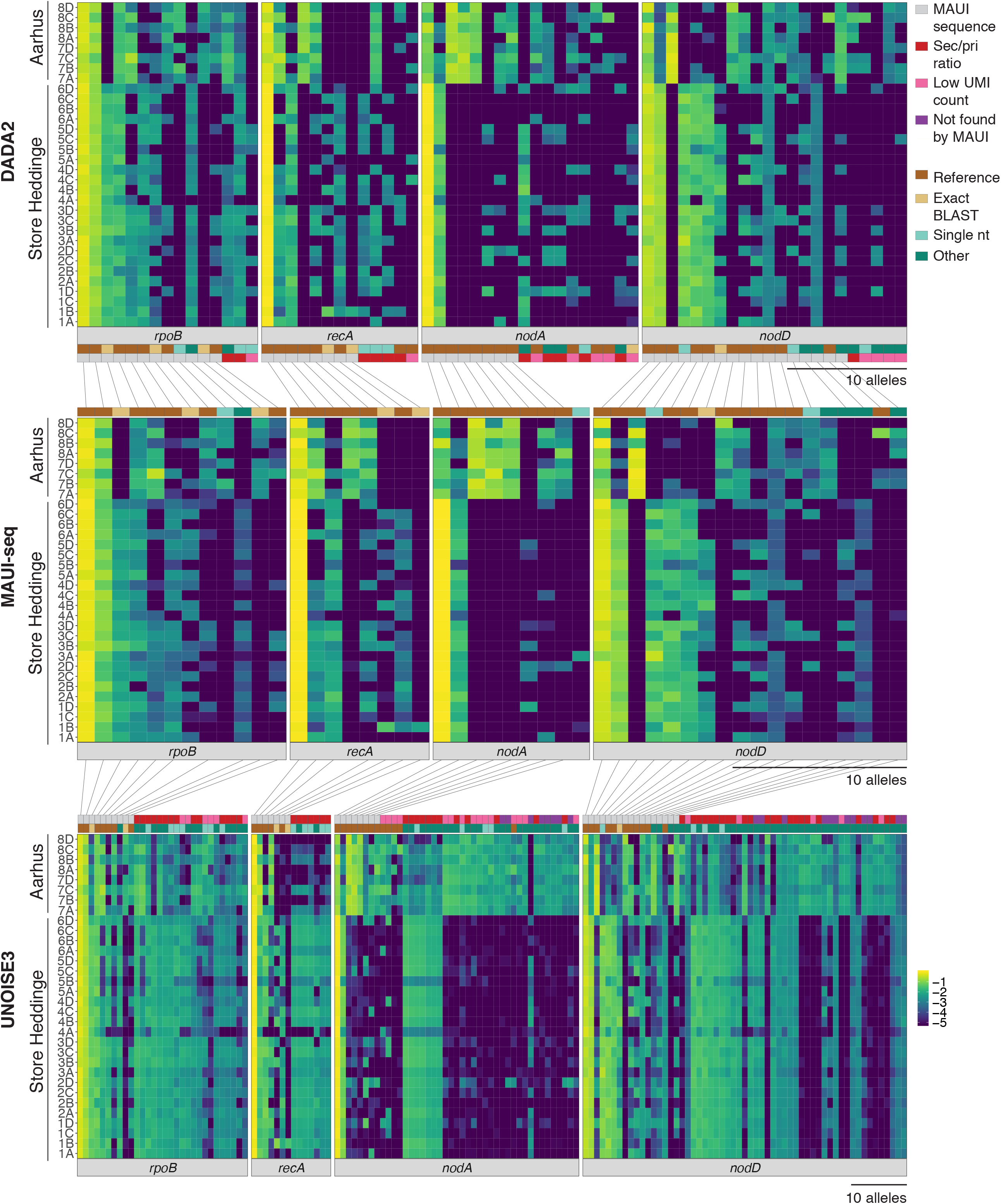
Amplicon diversity reported by MAUI-seq compared with the DADA2 and UNOISE3 analysis pipelines. Data are for four genes from nodule samples from two geographic locations, Store Heddinge (1-6) and Aarhus (7-8). Letters A-D denote the replicates within each plot (**Supplementary Figure 5**). Heatmap of the log10 transformed relative allele abundance of sequence clusters for individual genes. Lines connect identical sequences found by different clustering methods. Evidence that sequences are likely to be genuine is denoted by classifying them as **reference** (100% identity in at least 1 of 196 *Rhizobium leguminosarum* symbiovar *trifolii* genomes [29]), **exact BLAST** (100% query coverage and 100% identity against the whole-genome shotgun contigs BLAST database), **single nt** (one nt difference from either reference or exact BLAST match), and **other**. Sequences not reported by MAUI were classified as **sec/pri ratio** (rejected as erroneous because of a high secondary to primary ratio), **low UMI count** (not reported because too rare), **not found by MAUI** (no accepted UMIs).

The allele frequency distributions were different at Aarhus and Store Heddinge (**Figure 3**), and the two sites were clearly separated by the first principal component in a Principal Component analysis (PCA) for MAUI-seq, DADA2 and UNOISE3 sequences. (**Figure 4** and **Supplementary Figure S6-S8**). The amplicon sequencing has sufficient resolution to characterize geospatial variation in allele frequencies. For example, MAUI-seq, DADA2 and UNOISE3 can all clearly identify several highly abundant sequences from one location that are either absent or present in very low frequency in samples from the other location (**Figure 3**). To quantify the genetic differentiation between the Aarhus and Store Heddinge sites, we calculated fixation indices (*F*_ST_). Considering all four target genes combined, the MAUI-seq output resulted in the highest *F*_ST_ value followed by DADA2 and UNOISE3 (**Table 2, Figure 4** and **Supplementary Figure S9-S11**). For all individual genes, MAUI-seq also produced the highest *F*_ST_ estimates, and the differences were especially pronounced for *nodA*, which also showed the highest overall level of differentiation (**Table 2** and **Supplementary Figure S9-S11**). The lower genetic differentiation estimated based on DADA2 and UNOISE3 results, compared to those of MAUI-seq, reflects the inclusion of an increased number of erroneous sequences, which are less differentiated between the two sampled sites than the real sequences (**Figure 3**).

**Figure 4.**
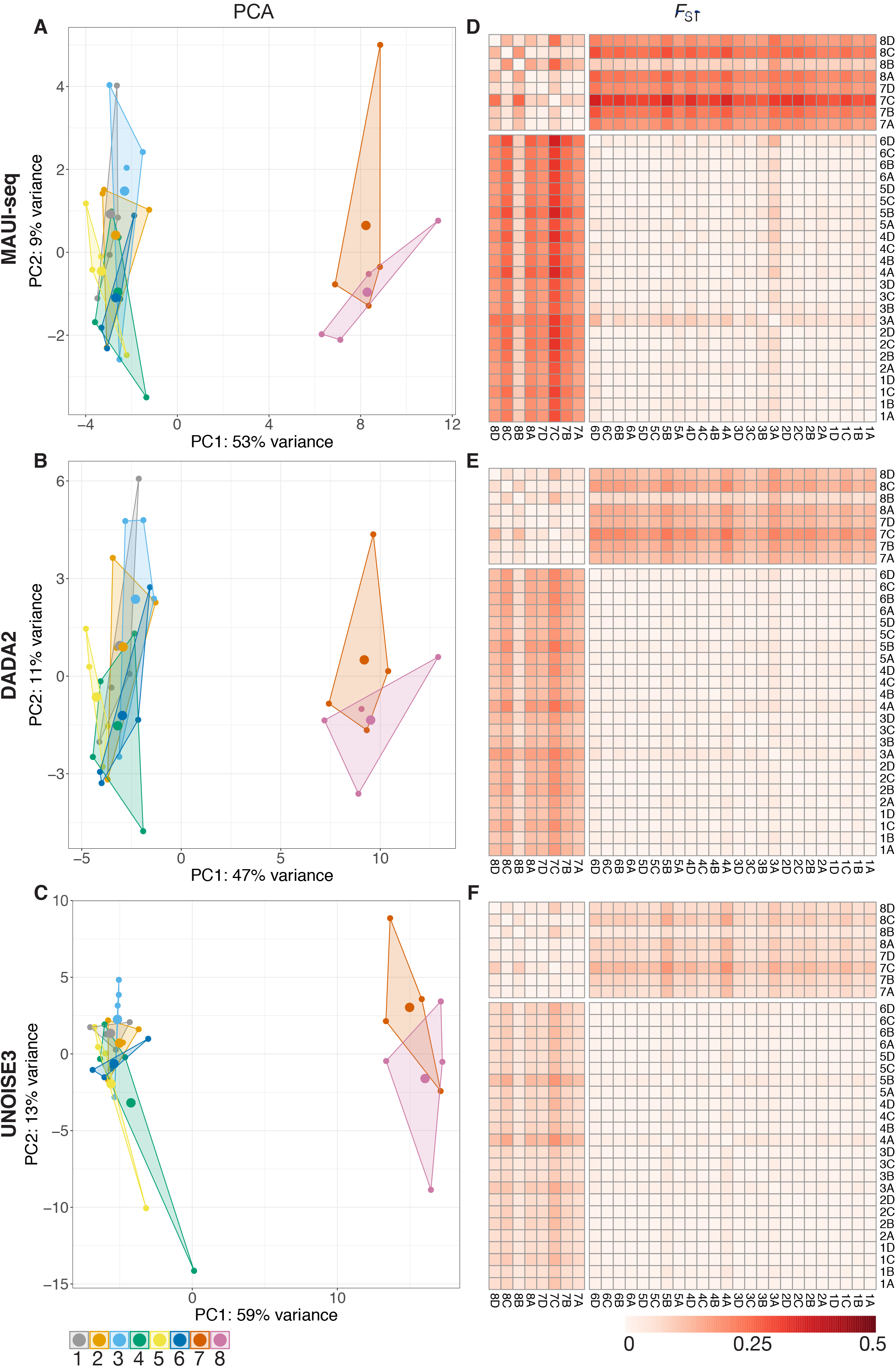
Genetic differentiation between populations visualised by Principal Component Analysis (**A-C**) and *F*_ST_ (**D-F**) of *Rlt* diversity in root nodule samples (8 sites, 4 replicates). Three analysis pipelines are compared: MAUI-seq (**A,D**), DADA2 (**B,E**), UNOISE3 (**C,F**). The PCA analysis was based on log10 transformed relative allele abundance. *F*_ST_ analysis was based on relative allele abundance. Data from all four genes (*rpoB*, *recA*, *nodA*, and *nodD*) were included in the analysis.

Since it was clear from the DNA mixture experiment that the choice of DNA polymerase could significantly affect error rates, we sampled root nodules from 13 additional clover field plots (the Field-Samples-2 dataset) and amplified each sample (a pool of one hundred root nodules) using Platinum and Phusion polymerases in parallel. For samples amplified using Platinum, MAUI-seq detected fewer sequences than DADA2 and UNOISE3 for the two core genes, but the same number of reference sequences were detected (**Table 3**). DADA2 included two chimeric sequences that were filtered out by MAUI-seq due to a high ratio of secondary to primary occurrences (**Additional File 2**). UNOISE3 detected twice as many sequences as DADA2 and MAUI-seq for the accessory genes, but most of the additional sequences had no associated UMIs and were classified as “other” (**Table 3, Additional File 2**). For samples amplified using Phusion, MAUI-seq and DADA2 detected a similar number of sequences (**Table 3**). All nine UNOISE3 *rpoB* sequences that were not accepted by either MAUI-seq or DADA2 (**Additional File 2**) are putative chimeric sequences with two parental sequences of higher abundance. For *nodA*, MAUI-seq includes three sequences that have a single nucleotide difference from a reference sequence, but all have a good ratio of secondary to primary reads, so we hypothesise that these are true sequences. Some reference or exact blast hit sequences were included by DADA2 but not by MAUI-seq because their abundance was estimated by DADA2 to be above the 0.001 threshold, but MAUI-seq estimated that they were rarer.

**Table 3.** The effect of polymerase choice. Total number of detected sequence clusters in root nodule samples (Field-Samples-2) amplified using Phusion (proofreading) or Platinum (non-proofreading) polymerases. Sequences were clustered using MAUI-seq, DADA2, and UNOISE3.

|  |  | Platinum |  |  | Phusion |  |  |
| --- | --- | --- | --- | --- | --- | --- | --- |
| Gene |  | MAUI-seq | DADA2 | UNOISE3 | MAUI-seq | DADA2 | UNOISE3 |
| *rpoB* | **Total** | **16** | **24** | **26** | **15** | **15** | **20** |
|  | **Reference\*** | 9 | 9 | 7 | 8 | 9 | 7 |
|  | **Exact** | 3 | 3 | 2 | 3 | 3 | 2 |
|  | **Single nt\*** | 3 | 7 | 8 | 3 | 2 | 5 |
|  | **Other\*** | 1 | 5 | 9 | 1 | 1 | 6 |
| *recA* | **Total** | **9** | **10** | **12** | **8** | **9** | **10** |
|  | **Reference** | 5 | 5 | 4 | 5 | 5 | 4 |
|  | **Exact** | 0 | 1 | 1 | 0 | 1 | 1 |
|  | **Single nt** | 3 | 3 | 3 | 3 | 2 | 3 |
|  | **Other** | 1 | 1 | 4 | 0 | 1 | 2 |
| *nodA* | **Total** | **18** | **14** | **35** | **17** | **11** | **34** |
|  | **Reference** | 7 | 10 | 8 | 9 | 9 | 9 |
|  | **Exact** | 0 | 1 | 0 | 0 | 0 | 0 |
|  | **Single nt** | 6 | 1 | 4 | 6 | 1 | 4 |
|  | **Other** | 5 | 2 | 22 | 2 | 1 | 21 |
| *nodD* | **Total** | **20** | **17** | **46** | **27** | **24** | **71** |
|  | **Reference** | 10 | 12 | 12 | 16 | 16 | 15 |
|  | **Exact** | 0 | 0 | 0 | 0 | 0 | 0 |
|  | **Single nt** | 6 | 3 | 6 | 5 | 4 | 6 |
|  | **Other** | 4 | 2 | 28 | 6 | 3 | 50 |

\* Output sequences were classified into **reference** (100% identity in at least 1 of 196 *Rhizobium leguminosarum* symbiovar *trifolii* genomes [29]), **exact BLAST** (100% query coverage and 100% identity against the whole-genome shotgun contigs BLAST database), **single nt** (one nt difference from either reference or exact BLAST match), and **other**.

Both MAUI-seq and DADA2 identify and remove sequences that appear to be errors (base substitutions or chimeras), but they use completely different evidence. As a result, they do not always make the same decision, as illustrated for a small set of representative data in **Table 4** (the *rpoB* sequences amplified by Phusion). While DADA2 examines the sequences and rejects those that are likely to be generated from more abundant sequences in the sample, MAUI-seq does not use the actual sequence but bases decisions on how frequently a sequence occurs as a secondary sequence with the same UMI as another (primary) sequence. Sequences ranked 5 and 6 (**Table 4**) are both potential chimeras of the more abundant sequences 1-4. Both DADA2 and MAUI-seq reject sequence 6 and accept sequence 5. Sequence 6 has a secondary/primary ratio of 103/118, which is above the default threshold of 0.7, so MAUI-seq rejects it as a likely error. On the other hand, the ratio for sequence 5 is 71/229. This is well below the threshold, but it is higher than other sequences with a similar primary count, e.g. sequence 9 (15/270). A possible explanation is that some of the reads for sequence 5 are generated as chimeras but others are genuine, since is entirely plausible that new alleles are generated by recombination between existing alleles. To some extent, MAUI-seq compensates for this because it allocates sequence 5 a relatively low count and hence lower ranking (8) than it has in the raw reads or the DADA2 analysis. There are two further sequences, 10 and 29, that are rejected by DADA2 as potential chimeras but accepted by MAUI-seq (**Additional file 2** Field-Samples-2-phusion-rpoB); in both cases they have secondary sequence counts well below the threshold, so MAUI-seq accepts them as genuine. DADA2 included an *rpoB* sequence that does not have any associated UMIs (sequence 41), and appears to be a chimera of two more abundant sequences (sequence 3/4/5 and sequence 11) (**Table 4**). MAUI-seq counts UMIs, not individual reads, and the default setting is to require that the primary sequence has at least two more reads than the next most frequent sequence (if any) that has the same UMI. This enriches for genuine sequences, which are generally more abundant than errors, but it means, of course, that the number of counts is much lower than the number of reads. In fact, for this particular set of data, the number of UMIs is orders of magnitude smaller than either the raw reads or the DADA2 count, although still sufficient to provide good estimates of the relative abundance of the sequences that make up the bulk of the population. The main reason for the low UMI count is that the number of reads per UMI was suboptimal in these data for the *rpoB* gene: only 18% of the UMIs had more than one read, and MAUI-seq discards single-read UMIs by default. By contrast, in the equivalent data for the *recA* gene in the same study (**Additional file 2** Field-Samples-2-phusion-recA), 37.5% of UMIs had more than one read, making more effective use of the available sequence reads.

**Table 4.** A comparison between DADA2 and MAUI-seq for a subset of the Field-Samples-2 data summarised in Table 3: the *rpoB* sequences from samples amplified by Phusion (proofreading) polymerase. Red cells refer to rejected sequences. Green cells refer to sequences, which are accepted by MAUI-seq, while DADA2 rejects them as potential chimeras. Yellow cells refer to sequences filtered out due to low UMI count by MAUI-seq.

| Raw reads |  | MAUI |  |  |  | DADA2 |  |  |
| --- | --- | --- | --- | --- | --- | --- | --- | --- |
| Rank | count | rank | UMI primary | UMI | accepted | rank | count | accepted |
| 1 | 99431 | 1 | 7459 | 197 | yes | 1 | 54758 | yes |
| 2 | 86751 | 2 | 7067 | 155 | yes | 2 | 48402 | yes |
| 3 | 70318 | 3 | 3668 | 95 | yes | 3 | 44412 | yes |
| 4 | 47337 | 4 | 1898 | 106 | yes | 4 | 28339 | yes |
| 5 | 13190 | 8 | 229 | 71 | yes | 5 | 7854 | yes |
| 6 | 11786 | 9 | 118 | 103 | no | none | NA | no |
| 7 | 10490 | 5 | 489 | 19 | yes | 6 | 6009 | yes |
| 8 | 9630 | 6 | 362 | 13 | yes | 7 | 5414 | yes |
| 9 | 4738 | 7 | 270 | 15 | yes | 8 | 2757 | yes |
| 10 | 4290 | 12 | 62 | 15 | yes | none | NA | no |
| 11 | 3223 | 11 | 90 | 3 | yes | 9 | 2041 | yes |
| 20 | 1950 | 10 | 96 | 6 | yes | 10 | 981 | yes |
| 29 | 1504 | 13 | 42 | 10 | yes | none | NA | no |
| 39 | 1063 | 14 | 35 | 2 | yes | 12 | 618 | yes |
| 41 | 946 | none | 0 | 0 |  | 11 | 721 | yes |
| 43 | 826 | 15 | 34 | 0 | yes | 13 | 434 | yes |
| 51 | 567 | 16 | 22 | 3 | yes | 14 | 341 | yes |
| 63 | 415 | 24 | 7 | 0 | (yes) | 15 | 208 | yes |

## Discussion

We propose a new HTAS method (MAUI-seq) designed to assess genetic diversity within or across species. It uses global UMI-based errors rates to detect potential PCR artefacts such as chimeras and single-base substitutions more robustly than the widely-used ASV clustering methods, DADA2 and UNOISE3. The approach is potentially applicable to any study of amplicon diversity, including community diversity estimates based on 16S rRNA and other metabarcoding surveys using environmental DNA.

### Using UMIs to filter out chimeras and other errors

In the MAUI-seq approach, UMIs are used to reduce errors in two distinct ways. Since all reads with the same UMI should, in principle, be derived from the same initial template copy, any variation among them reflects errors. In some implementations, a consensus sequence is calculated [24], but we adopt the simpler approach of accepting the most abundant sequence, which will usually give the same result. Requiring more than one identical read before accepting a UMI creates an important quality filter that greatly reduces the number of rare (and usually erroneous) sequences, but as more reads are required, an increasing number of the original reads are discarded and the number of accepted counts declines. To strike a balance between quantity and quality, we chose to count a sequence provided it had at least two more reads than the next most frequent sequence with the same UMI, but this threshold could be adjusted if, for example, a markedly larger number of reads were available.

While the most abundant sequence associated with a UMI will usually be the correct one, it will sometimes happen that an erroneous sequence will predominate among the small number of reads actually sequenced, leading to these sequences being included among the recorded counts. These errors can be detected, though, by aggregating information across the whole set of samples. When a UMI is associated with more than one sequence, the secondary sequences are most often erroneous, so sequences that are relatively more abundant as secondary sequences than as the primary sequences associated with UMIs are likely to be erroneous. We recorded the number of times each sequence was found as the second sequence associated with a UMI, and found empirically that a suitable threshold for accepting sequences as genuine was that they occurred less than 0.7 times as often as secondary sequences as they occurred as primary sequences. This threshold can, however, be adjusted to reflect the error distribution observed in a particular study. We found that this approach was very effective in identifying known errors, particularly chimeras, which were generally the most abundant errors. Chimeras were rejected more effectively by MAUI-seq than by the two established ASV clustering methods, DADA2 and UNOISE3. Both of these rely on *de novo* rejection of sequences that could be constructed as recombinants of other sequences that are more abundant in the sample [13]. This method risks rejecting sequences that appear to be recombinant but are genuine alleles, which may not be uncommon, particularly in intraspecific samples. Our approach, by contrast, uses information on the observed error rates in the data (detected using UMIs) to decide whether a sequence is likely to be genuine, regardless of its actual sequence and relationship to other sequences. Sequences that could be generated as chimeras, or that differ by a single nucleotide from a more abundant sequence, may be accepted as genuine if they are more abundant than expected from their rate of occurrence as minor sequences associated with UMIs. In our study, this approach eliminated many known errors and substantially improved our confidence in the remaining data, providing a powerful additional reason for using UMIs in metabarcoding studies of all kinds. While we found that a simple empirical threshold was effective, we noticed that the proportion of secondary sequences varied markedly across studies and genes, suggesting that an adjustable threshold might give further improvement. A useful future development might be to use the abundance of minor sequences associated with UMIs to generate a statistical model of error processes that would provide a firmer theoretical basis for the classification of sequences.

### Using UMIs to reduce amplification bias

One motivation for the use of UMIs is to obtain more accurate relative abundance data by eliminating possible sequence-specific bias in the PCR amplification, which may be introduced by variation in polymerase and primer affinity for some DNA templates. Indeed, we observed that the Platinum polymerase preferentially amplified the SM170C *rpoB* allele, whereas the Phusion enzyme did not have this bias (**Table 1** and **Supplementary Figure S1A-C**). Allele variant bias was also shown for other target genes, although the ranking of the two enzymes was not always the same (**Table 1** and **Supplementary Figures S1-S4**). However, in our study, the use of UMIs did not correct the allele bias. This suggests that the bias was present in the initial round of copying using the target-specific primer, rather than in the subsequent amplification rounds. For our case study, at least, the choice of polymerase was much more important for accurate relative abundance data than the use of UMIs. The main advantage of UMIs was, rather, the ability to remove most sequencing errors, as discussed in the preceding section.

### Advantages of multiplexing several amplicons

Increasing the number of monitored amplicons to four increased our ability to robustly distinguish samples from two locations (**Figure 3-4** and **Supplementary Figure S6-S11**). Multiplexing could be used in other ways, for example to monitor several organisms in the same environment, or to increase read coverage profiling of single genetic markers such as 16S [30]. In addition, there is a technical benefit in sequencing multiple different targets together, because a lack of sequence diversity can cause Illumina base-calling issues [31].

### Optimization of the protocol

As with any metabarcoding project, the first important step is to design the primers carefully to amplify the entire target community with minimum bias, and we used a large database of known gene sequences to achieve this. Another consideration that is shared with other approaches is the choice of polymerase for PCR. For the samples studied here, with abundant template DNA, the proofreading enzyme was clearly superior in performance, although more costly. On the other hand, this enzyme may provide less robust amplification when the template is weak, as we have observed in another project aimed at rhizobial DNA in soil [32]. The use of UMIs introduces other design considerations. We used twelve random nucleotides (with some constraints), giving over four million potential UMI sequences, which was sufficient for the scale of our studies, but it would be simple to increase the UMI length if greater sequencing depth was planned. In any metabarcoding study, the choice of sequencing depth is, to some degree, made blindly because the diversity of templates is not known in advance, but UMI-based approaches need greater depth because it is UMIs that are counted, not reads, and the aim is to have several reads per UMI. There are many factors that affect the average number of reads per UMI, but our study is encouraging in that, without separate optimization, all of our target genes in all of our samples gave usable data. In fact, the number of reads per UMI were suboptimal in most cases. Given a fixed sequencing effort, reads per UMI could, if necessary, be increased by reducing the concentration of the forward UMI-bearing primer and/or of the sample DNA so that fewer distinct UMIs were initiated. With our parameters, at least two reads are needed before a UMI is counted, and a sufficient fraction of the UMIs need at least four reads so that some will have a secondary sequence as well as the primary sequence (with at least two reads more than the secondary).

### Future directions for MAUI-seq

HTAS is a valuable and widely-used approach for the study of microbial community diversity, but handling erroneous sequences introduced by the amplification and sequencing procedures has always been challenging. The use of UMIs allows MAUI-seq to greatly reduce the incidence of errors through two mechanisms. Firstly, the requirement that a UMI is associated with at least two identical reads eliminates many rare sequences that are predominantly erroneous. Secondly, sequences that are frequently generated as errors can be identified and removed because they occur unexpectedly often as minor components associated with UMIs that are assigned to more abundant sequences. These mechanisms are independent of any reference database and can recognise and retain genuine alleles that differ by a single nucleotide or match a potential chimera. This makes MAUI-seq particularly suited to studies of intraspecific variation, where the range of sequence divergence may be limited and not fully known in advance. However, the efficient elimination of erroneous sequences is also important in community studies such as those based on widely-used 16S primers, and MAUI-seq should be readily adaptable to this field. The analysis pipeline is very fast because no sequence alignment or database searching is involved; only the accepted final sequences would need to be characterised by comparison to a reference database.

Most HTAS studies report the relative proportions of the taxa in a community, but it would sometimes be valuable to estimate the absolute abundance of the microbes in the environmental sample. UMIs can potentially provide such information, if the initial template copying is carefully controlled so that the total number of distinct UMIs reflects the number of templates [26, 33]. While this would necessitate some additional steps at the start of the experimental protocol, it should still be possible to analyse the resulting sequences using the error-removal approaches provided by MAUI-seq. Alternatively, absolute abundance can be estimated by adding a spike of a known quantity of a recognisable target sequence to the sample before processing [11, 34, 35].

The addition of a UMI shortens the maximum length of target sequence that can be read, and the counting of UMIs rather than reads requires a higher depth of sequencing, but these limitations are increasingly unimportant as improvements in sequencing technology lead to increasing length, enabling long-read amplicon sequencing [36, 37], and numbers of reads. As implemented in MAUI-seq, UMIs are very effective in reducing the errors inherent in HTAS, and have the potential to improve the quality of any amplicon-based study of diversity.

## Materials and methods

### Preparation of DNA mixtures

Two *Rlt* strains (SM3 and SM170C) were chosen based on their *recA*, *rpoB*, *nodA*, and *nodD* sequence divergence, with a minimum of 3 base pair differences in the amplicon region required for each gene. Strains were grown on Tryptone Yeast agar (28°C, 48hrs). Culture was resuspended in 750ul of the DNeasy Powerlyzer PowerSoil DNA isolation kit (QIAGEN, USA) and DNA was extracted following the manufacturer’s instructions. DNA sample concentrations were calculated using QuBit (Thermofisher Scientific Inc., USA). DNA samples of the two strains were diluted to the same concentration and mixed in various ratios (**Supplementary Table S1)**.

### Preparation of environmental samples

For Field-Samples-1 data, white clover (*Trifolium repens*) root nodules were collected from two locations: Store Heddinge, Denmark (6 plots) and Aarhus University Science Park, Aarhus, Denmark (2 plots) (**Supplementary Figure S4**). The clover varieties sampled were Klondike (Store Heddinge) and wild white clover, (Aarhus). 100 large pink nodules were collected from 4 points on each plot, making a total of 32 samples. Nodules were stored at −20°C until DNA extraction. Nodule samples were thawed at room temperature and crushed using a sterile homogeniser stick. Crushed nodules were mixed with 750μl Bead Solution from the DNeasy PowerLyzer PowerSoil DNA isolation kit (QIAGEN, USA) and DNA was extracted following the manufacturer’s instructions. DNA sample concentrations were measured using a Nanodrop 3300 instrument (Thermofisher Scientific Inc., USA).

For Field-Samples-2 data, root nodules were additionally sampled from 13 white clover conventionally-managed field trial plots at Store Heddinge, Denmark (Sample 1A-13A, **Additional File 2**). All plots were sown under the same conditions in 2017. Three to ten clover plants were sampled from one point in each plot and the 100 largest nodules collected. Nodules were stored at −20°C, and DNA was extracted for each sample using the Qiagen DNeasy PowerLyzer PowerSoil DNA isolation kit, as above. Samples were processed independently with Platinum (non-proofreading) and Phusion (proofreading) polymerases to evaluate the method dependency on polymerase choice, as described in the following sections.

### PCR and purification

Primer sequences were designed for two *Rlt* housekeeping genes, recombinase A (*recA*) and RNA polymerase B (*rpoB*) and for two *Rlt* specific symbiosis genes, *nodA* and *nodD* (**Additional File 1: Table S1**).

The three primers are a target-gene forward inner primer, a universal forward outer primer, and a target-gene reverse primer. The concentration of the inner forward primer was 100-fold lower than the universal forward outer primer and the reverse primer (**Figure 1**) in order to reduce the competitiveness of this primer compared to the outer primer. The inner primer is essential for the first round of amplification, but its participation is undesirable in later rounds as it would assign a new unique UMI to an existing amplicon. The PCR reaction mixture and thermocycler programme are provided (**Additional File 1: Tables S2 and S3**).

PCRs were undertaken individually for each primer set using Platinum Taq DNA polymerase (Thermofisher Scientific Inc., USA) (**Additional File 1: Table S2**) and subsequently pooled and purified using AMPure XP Beads following the manufacturer’s instructions (Beckman Coulter, USA). Successful PCR amplification was confirmed by running a 0.5X TBE 2% agarose gel at 90V for 2 hours.

For the DNA mixture samples, PCRs were run in triplicate. DNA from single strains was also processed as a control to determine the level of cross contamination between samples. Some samples were also amplified using Phusion High-Fidelity polymerase (Thermofisher Scientific Inc., USA), to evaluate whether use of a proof-reading polymerase improved the quality of the results using the PCR program described in **Additional File 1: Table S2** and **Table S4**.

### Nextera indexing for multiplexing and MiSeq sequencing

Samples were indexed for multiplexed sequencing libraries with Nextera XT DNA Library Preparation Kit v2 set A (Illumina, USA) using the Phusion High-Fidelity DNA polymerase (Thermofisher Scientific Inc., USA). PCR reaction mixture and programme are detailed in **Additional File 1: Tables S6 and S7** Indices were added in unique combinations as specified in the manufacturer’s instructions (Illumina, USA).

The PCR product was purified on a 0.5X TBE 1.5% agarose gel and extracted with the QIAQuick gel extraction kit (QIAGEN, USA) (expected band length: ~454bp). PCR amplicon concentrations were quantified using GelAnalyzer2010a and normalised to 10nM [38]. A pooled sample was quantified and checked for quality by Bioanalyzer (Agilent, USA) before sequencing using Illumina MiSeq (2×300bp paired end reads) by the University of York Technology Facility. A detailed protocol is available in **Additional File 1**.

### Read processing and data analysis

The PEAR assembler was used to merge paired ends [39]. Python scripts were used to separate the merged reads by gene (MAUIsortgenes.py) and to calculate allele frequencies both with and without the use of UMIs (MAUIcount.py). The scripts are available in the GitHub repository https://github.com/jpwyoung/MAUI. Sequences were clustered by UMI, and the number of unique UMIs was counted for each distinct sequence, provided that sequence had at least two more reads with that UMI than any other sequence. In cases where two or more sequences were associated with the same UMI, the second most abundant sequence was noted, and sequences that occurred more than 0.7 times as often as second sequences than as the main sequence associated with a UMI were filtered out of the results as putative PCR-induced chimeras or other errors. Sequences with primers removed (ignoring UMIs) were also clustered using DADA2 (version 1.8) [19] and UNOISE3 (USEARCH version 11.0.667) [21] with default settings. An overall read frequency filter of 0.1% was applied to DADA2 and UNOISE3 outputs to match MAUI-seq accepted sequences filtering. Scripts used for DADA2, UNOISE3, and figure generation are available in **Additional file 3**, **4**, and **5**, respectively. Output abundance data were then processed for statistical analysis and figure generation using various R packages (**Additional File 3, 4,** and **5**; [40, 41]). Principal components were calculated with the R ‘prcomp’ package using singular value decomposition to explain the *Rhizobium* diversity and abundance within each sub-plot sample. Differences in allele frequencies between samples were quantified using Bray-Curtis beta-diversity estimation using the R package ‘vegdist.’ PERMANOVA tests were performed using the R package ‘adonis’. Empirical Bayes estimator of *F*_ST_ was calculated using the R package ‘FinePop’.

## Acknowledgements

We thank David Sherlock for his experimental expertise in developing this method, the University of York Technology Facility for sequencing, Simon Kelly for DADA2 expertise, Asger Bachmann, Terry Mun, Maria Izabel A. Cavassim, and Marni Tausen for preliminary data analysis and script development, and DLF for access to their clover field trials. This work was funded by grant no. 4105-00007A from Innovation Fund Denmark (S.U.A.). Initial development of the method was funded by the EU FP7-KBBE project LEGATO (J.P.W.Y).

## Author contributions

Conceptualization: JPWY; Methodology: JPWY, SUA; Software: BF, SM, JPWY; Validation: BF, SM, JPWY, SUA; Formal analysis: BF, SM, JPWY; Investigation: BF, SM; Resources: JPWY, SUA, VPF; Data curation: BF, SM, JPWY; Writing - original draft: BF, SM; Writing - review and editing: BF, SM, JPWY, SUA, VPF; Visualisation: BF, SM; Supervision: JPWY, SUA, VPF; Project administration: JPWY, SUA; Funding acquisition: JPWY, SUA.

## Availability of supporting data

Raw Illumina reads are available in the SRA repositories with accession numbers [SRP221010 (Synthetic mix and Field-Samples-1) and SRP238323 (Field-Samples-2)]. MAUI-seq scripts are available in the GitHub repository https://github.com/jpwyoung/MAUI. A detailed protocol for sampling, sample preparation, and read processing is available in **Additional file 1**. Scripts used for DADA2, UNOISE3, and figure generation are available in **Additional file 3**, **4**, and **5**, respectively. Detailed output sequences for all three methods are available in **Additional file 2**.

## Ethics approval and consent to participate

Not applicable

## Consent for publication

Not applicable

